# Antibodies targeting Crimean-Congo hemorrhagic fever virus GP38 limit vascular leak and viral spread

**DOI:** 10.1101/2024.05.23.595578

**Authors:** Felix Pahmeier, Stephanie R. Monticelli, Xinyi Feng, Christy K. Hjorth, Albert Wang, Ana I. Kuehne, Russell R. Bakken, Thomas G. Batchelor, Saeyoung E. Lee, Marissa Middlecamp, Lauren Stuart, Dafna M. Abelson, Jason S. McLellan, Scott B. Biering, Andrew S. Herbert, Kartik Chandran, Eva Harris

## Abstract

Crimean-Congo hemorrhagic fever virus (CCHFV) is a priority pathogen transmitted by tick bites, with no vaccines or specific therapeutics approved to date. Severe disease manifestations include hemorrhage, endothelial dysfunction, and multiorgan failure. Infected cells secrete the viral glycoprotein GP38, whose extracellular function is presently unknown. GP38 is considered an important target for vaccine and therapeutic design as GP38-specific antibodies can protect against severe disease in animal models, albeit through a currently unknown mechanism of action. Here, we show that GP38 induces endothelial barrier dysfunction *in vitro*, and that CCHFV infection, and GP38 alone, can trigger vascular leak in a mouse model. Protective antibodies that recognize specific antigenic sites on GP38, but not a protective neutralizing antibody binding the structural protein Gc, potently inhibit endothelial hyperpermeability *in vitro* and vascular leak *in vivo* during CCHFV infection. This work uncovers a function of the secreted viral protein GP38 as a viral toxin in CCHFV pathogenesis and elucidates the mode of action of non-neutralizing GP38-specific antibodies.

## Introduction

The World Health Organization has designated Crimean-Congo hemorrhagic fever virus (CCHFV) a priority pathogen due to its high lethality and lack of effective countermeasures. CCHFV belongs to the family *Nairoviridae* in the order *Bunyavirales* and contains a tri-segmented negative-strand and ambisense genome. Transmission to humans via tick bite is followed by an incubation period of 5-13 days, leading to initial nonspecific symptoms such as fever and malaise *(1)*. In severe cases, progression to endothelial barrier dysfunction, shock syndrome, and multi-organ failure, including liver and spleen pathology, results in an overall case fatality rate of 5-40% *(1–4)*. The triggers of this pathology and disease progression are not well understood but are generally attributed to uncontrolled viral replication and the release of pro-inflammatory cytokines *(2)*. Further, the characteristics of a protective immune response have not been fully elucidated, although a role for protective antibodies has been implicated in leading to milder disease manifestations *(5, 6)*. Recent studies of human survivor cohorts found that in addition to the presence of virus-binding, neutralizing antibodies targeting the structural glycoprotein Gc, antibodies binding to GP38, a secreted glycoprotein generated by cleavage of the viral glycoprotein precursor complex, are also elicited *(7–10)*. GP38-specific monoclonal antibodies (mAbs), as well as GP38-based vaccines, are protective in murine models of CCHFV challenge, albeit via an unknown mechanism of action that is independent of neutralization and Fc-dependent functions *(8, 9, 11–15)*. Although intracellular GP38 has been shown to play a role in virion assembly in the secretory pathway *(16)*, the specific functions of the extracellular form of the protein are unknown.

These characteristics are highly reminiscent of the non-structural protein 1 (NS1) of flaviviruses, a glycoprotein involved in intracellular replication and assembly whose secreted form has been implicated in viral pathogenesis and shown to be a target of non-neutralizing protective antibodies *(17–20)*. NS1 can trigger endothelial barrier dysfunction and vascular leak independent of viral infection through an endothelial cell-intrinsic pathway *(20–22)*, and NS1-specific mAbs can prevent induction of endothelial dysfunction and vascular leak as well as protect against lethal dengue virus (DENV) infection in a mouse model *(17–19)*.

In this study, we show that GP38 can induce vascular leak and endothelial barrier dysfunction in CCHFV infection through an EC-intrinsic pathway, presenting a novel function of this secreted glycoprotein as a viral toxin. Furthermore, our data demonstrates that GP38-targeting mAbs protect mice from vascular leak and viral dissemination, uncovering a previously unrecognized mechanism of action of these protective mAbs.

## Results

### CCHFV infection causes vascular leak and endothelial barrier dysfunction

Building on the similarities of CCHFV GP38 with flavivirus NS1, we hypothesized that GP38 may have a similar function as a secreted viral toxin by triggering endothelial barrier dysfunction and vascular leak during CCHFV infection (Fig. 1A). To test this, we used a transient immunosuppression-based murine model of CCHFV infection to investigate vascular leak during infection *(11, 12, 18, 23, 24)*. C57BL/6 mice were infected with 100 plaque-forming units (PFU) of CCHFV (strain IbAr10200) and subsequently treated with an anti-interferon alpha/beta receptor (IFNAR) mAb (clone MAR1-5A3) 24 hours post-exposure. Three days following infection, at the height of disease, a combination of tracer dyes (10 kDa-dextran conjugated to Alexa Fluor 680; Evans Blue) was intravenously injected to measure vascular leak. After dye circulation, whole blood (for serum) and tissues (liver, spleen, and kidney) were collected, and viral load and dye extravasation were assessed. High viral load was measured in the serum as well as in the liver, spleen, and kidney, indicative of extensive viral dissemination into distal tissues (Fig 1B). Measurement of the tracer dye in the liver, the major site of viral replication and pathology *(3, 4)*, revealed a significant increase in extravasated dyes compared to mock-infected mice, with distinct foci of vascular leak (Fig. 1C-D, fig. S1). Hypothesizing that GP38 may function as a secreted viral toxin, we further quantified levels of circulating GP38 by developing a quantitative sandwich ELISA utilizing two recently described GP38-specific mAbs that bind to distinct non-competitive epitopes on the protein *(9)* (fig. S2). Levels of GP38 in the bloodstream reached a mean concentration of 1 µg/mL, with a maximum of 5 µg/mL (Fig. 1E). These data demonstrate that vascular leak occurs during authentic CCHFV infection and establishes an *in vivo* model to measure the induction of vascular leak. Furthermore, these data show that GP38 circulates in the bloodstream of infected mice at appreciable levels.

**Figure 1.**
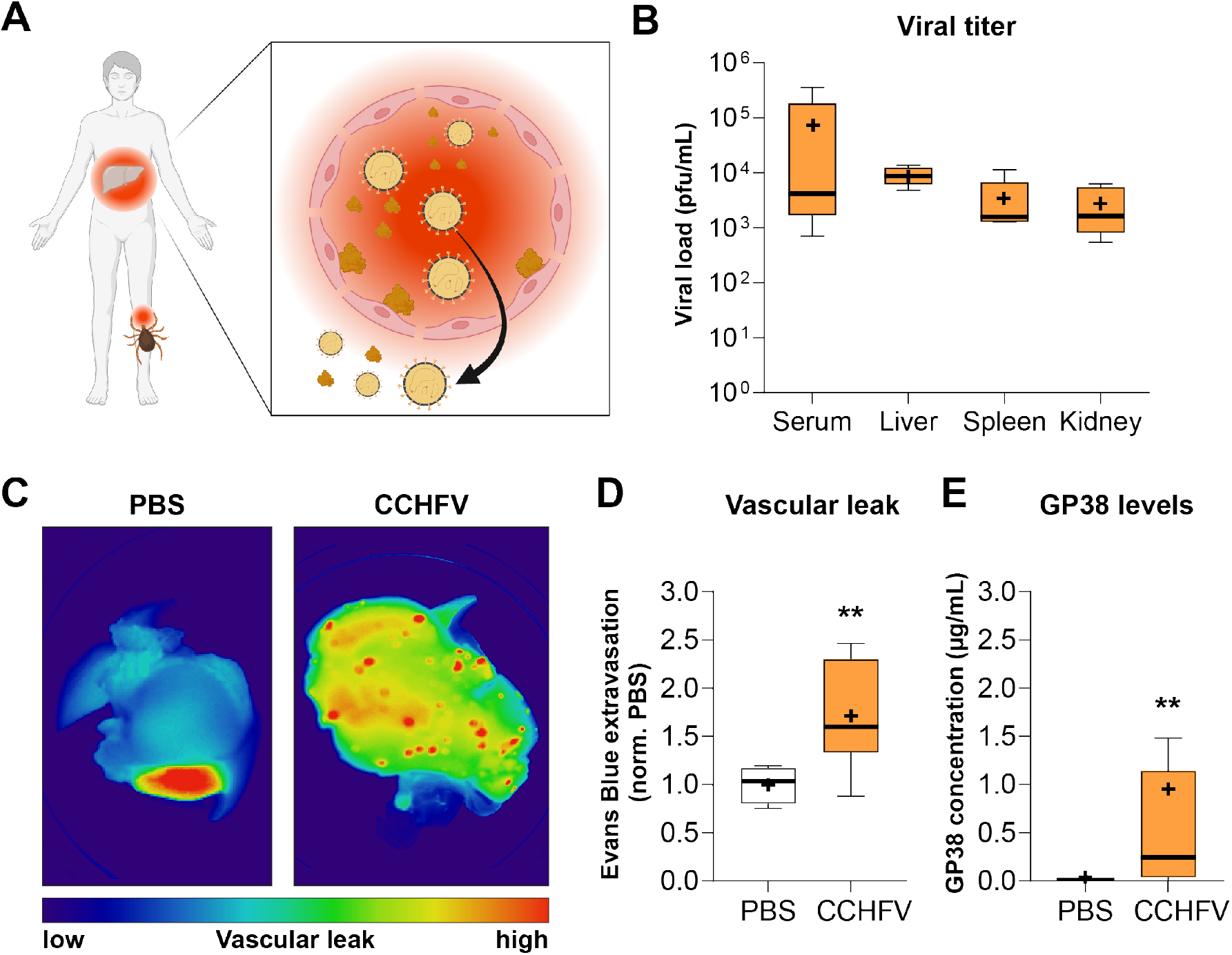
CCHFV infection leads to viral dissemination, vascular leak, and circulation of GP38. (A) Transmission of CCHFV by tick bites is followed by virus amplification in the bloodstream, endothelial dysfunction, and liver pathology. (B) Viral load in tissues (liver, spleen, kidney) and serum from mice infected with CCHFV and treated with anti-IFNAR mAb MAR1-5A3 three days post-infection (n=5-10). (C) Representative images of leak of tracer dye Dextran-680 in the liver of PBS-treated or CCHFV-infected mice. (D) Extravasation of tracer dye Evans Blue dye into the liver of PBS-treated or CCHFV-infected mice relative to PBS control animals (n=5-10). (E) Concentration of GP38 in the serum of PBS-treated or CCHFV-infected mice determined by quantitative sandwich ELISA (n=5-14). Data shown as a Tukey distribution with the median as a bar and mean as +. Statistical comparisons were performed by Mann-Whitney test with **, p < 0.01.

### CCHFV GP38 causes vascular leak independently of viral infection

The parallels with flavivirus NS1 and the observation of circulating GP38 led us to investigate whether GP38 alone is sufficient to trigger vascular leak in the absence of infection. We first tested the capacity of recombinant GP38 (fig. S3A-B) to cause vascular leak in an established dermal vascular leak model *(17, 21, 24)*. GP38, alongside negative and positive controls (PBS and DENV NS1, respectively) was injected intradermally in distinct spots in the dorsal dermis before intravenous administration of Dextran-680. Scanning of the dissected dermis showed a dose-dependent increase in tracer dye extravasation upon intradermal injection of 5 to 15 µg of GP38, similar to that observed for the positive control, DENV NS1 (Fig. 2A-B). Next, GP38 was administered into the bloodstream of mice, and induction of vascular leak was measured using the tracer dye-based systemic vascular leak assays as described above *(20, 24)*. Two days following retro-orbital intravenous injection of GP38, we found that systemic administration of GP38 caused an increase in tracer dye extravasation (Evans Blue) in the liver and spleen, but not in the kidney or lung, relative to an ovalbumin control (Fig. 2C-F). In summary, GP38 can cause vascular leak independently of viral infection, with a tropism comparable to the sites of pathology observed during natural viral infection *(3, 4)*.

**Figure 2.**
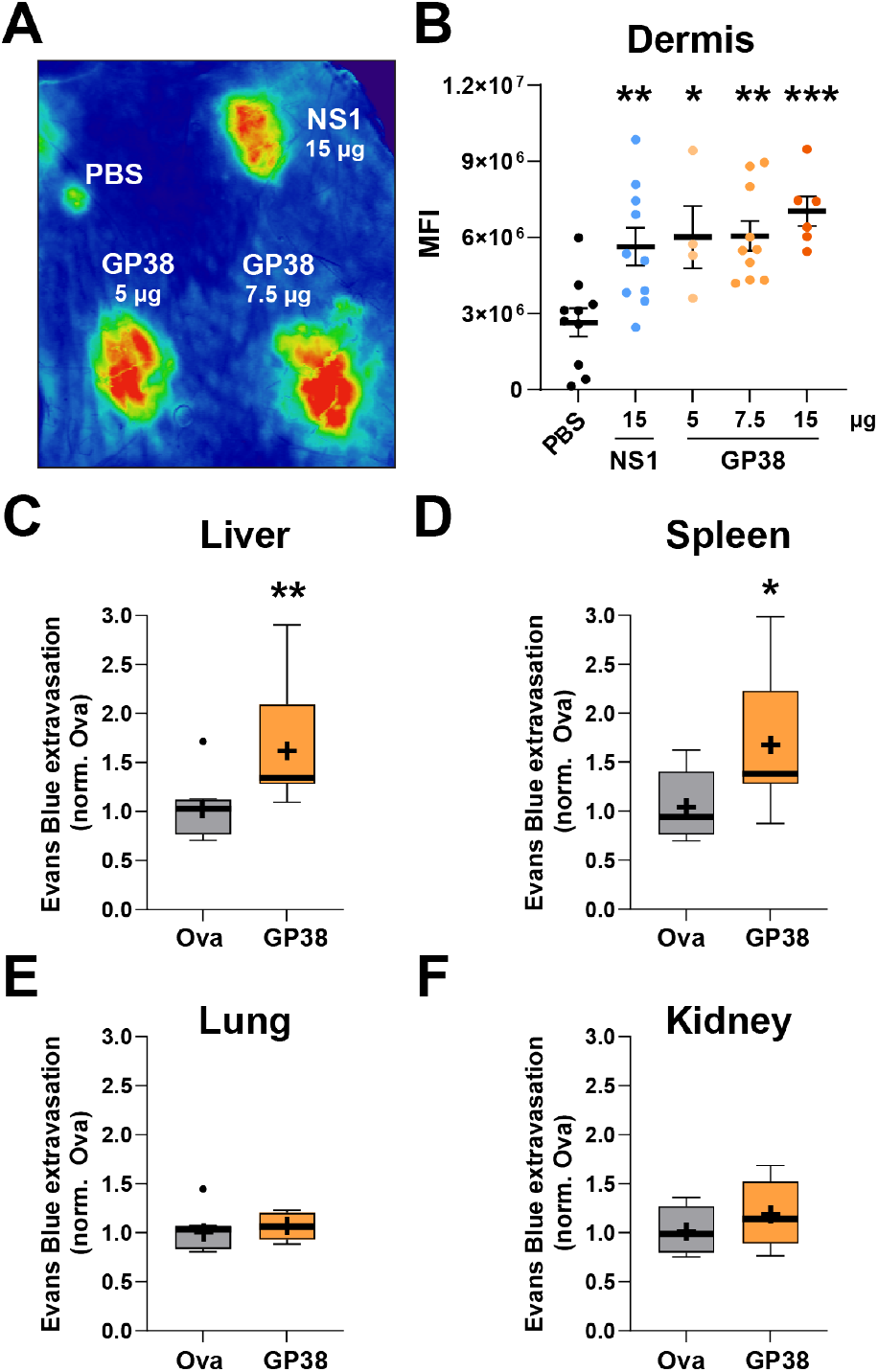
CCHFV GP38 causes vascular leak independently of virus infection. (A) Leak of the tracer dye Dextran-680 in the mouse dorsal dermis was measured after intradermal injection of CCHFV GP38 with the indicated controls (negative, PBS; positive dengue virus NS1) into the shaved backs of mice. One representative scan is shown. (B) Quantification of (A) as mean fluorescence intensity (MFI). (C-F) Mice were intravenously injected with 0.1 mg of GP38 or ovalbumin (Ova) and after two days, mice were intravenously injected with Evans Blue dye before organ harvest. Relative levels of Evans Blue dye extracted from the liver (C), spleen (D), lung (E) and kidney (F) compared to the ovalbumin control are shown as a Tukey distribution with the median as a bar and mean as + (n=8). Statistical comparisons were performed by Ordinary one-way ANOVA with Dunnett’s multiple comparison’s test (A) or Mann-Whitney tests (C-D) with *, p < 0.05; **, p < 0.01; ***, p < 0.001.

### Endothelial hyperpermeability and endothelial glycocalyx layer disruption is triggered by CCHFV GP38

We next investigated potential mechanisms underlying the observed GP38-mediated vascular leak. Endothelial dysfunction has been reported in human CCHFV infection, and GP38 is secreted from infected cells and circulates in the bloodstream *(25, 26)* (Fig. 1E), which led us to hypothesize that GP38 might interact with the endothelial cells lining the bloodstream and trigger endothelial dysfunction (Fig. 1A), similar to what has been shown for flavivirus NS1 *(22)*. Endothelial barrier dysfunction can be tested *in vitro* by measuring transendothelial electrical resistance (TEER) and integrity of the endothelial glycocalyx layer (EGL) (Fig. 3A). Human pulmonary microvascular endothelial cells (HPMECs) treated with 2.5 µg/mL GP38, as well as HPMEC treated with positive controls tumor necrosis factor (TNF)-α (10 ng/mL) and DENV NS1 (10 µg/mL), exhibited a reduction in TEER six hours post-treatment (hpt), indicative of endothelial hyperpermeability (Fig. 3B-C). Importantly, the structural CCHFV glycoprotein Gc did not trigger hyperpermeability (Fig. 3B-D). We next tested the amounts of GP38 required to cause endothelial barrier dysfunction and observed a dose-dependent increase in endothelial hyperpermeability in HPMEC down to concentrations of 0.25 µg/mL (Fig. 3D, Supp. Fig. 4A), approximately 20-fold lower than the maximum concentration of circulating GP38 measured during CCHFV infection *in vivo* (Fig. 1E). We further tested whether GP38 could trigger hyperpermeability in human skin-derived microvascular endothelial cells (HMEC), as the skin is the site of CCHFV transmission during a tick bite, and we found a comparable dose-dependent induction of hyperpermeability (Fig. 3E, fig. S4B). To ascertain specificity, we complexed GP38 with the protective GP38-specific mAb 13G8 *(13, 14)* and found that 13G8, but not the NS1-specific isotype control 2B7 *(17)*, was able to block GP38-triggered endothelial hyperpermeability (Fig. 3F).

**Figure 3.**
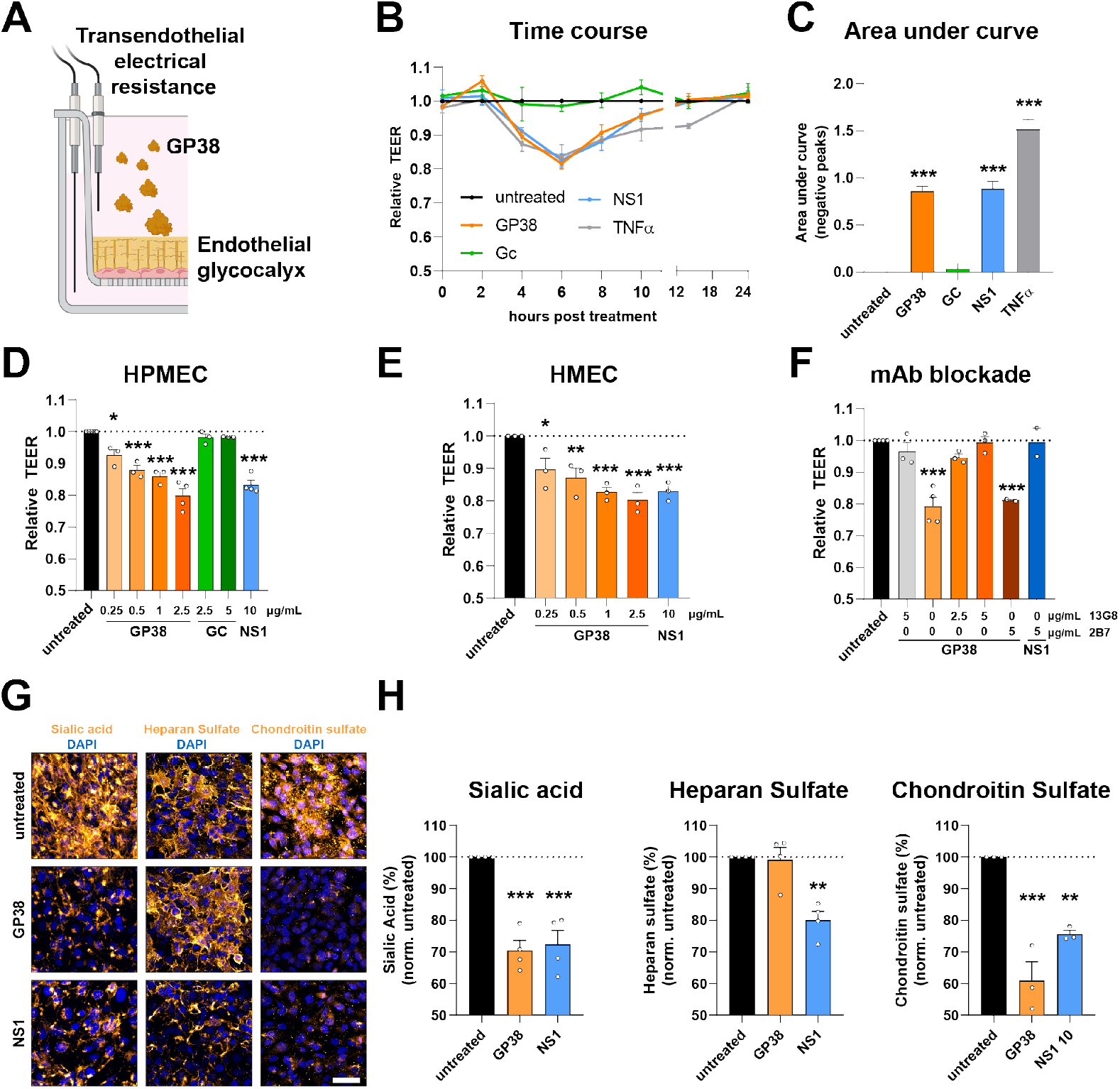
CCHFV GP38 triggers endothelial hyperpermeability and endothelial glycocalyx layer component disruption. (A) Endothelial barrier dysfunction can be assessed by measuring transendothelial electrical resistance (TEER) in a Transwell system and staining of endothelial glycocalyx layer (EGL) components. (B) Relative TEER of human pulmonary microvascular endothelial cells (HPMEC) treated with GP38 or PBS and measured over a 24-hour time-course (n=3). (C) Quantification of (B) by area under the curve of the negative peaks compared to the untreated control (n=3). (D-F) Relative TEER of a monolayer of HPMEC (D, F) and human dermal microvascular endothelial cells (HMEC) (E) at 6 hours post-treatment (hpt) with GP38 complexed or not with mAb (n≥2), normalized to the untreated control. (G-H) HPMEC were stained for EGL components sialic acid, heparan sulfate, and chondroitin sulfate 6 hours after GP38 or dengue virus NS1 treatment. Representative images are shown in (G) (scale bar = 50 µm), and MFI was quantified and normalized to the untreated control in (H) (n≥3). Statistical comparisons were performed by Ordinary one-way ANOVA with Holm-Sidak test with *, p < 0.05; **, p < 0.01; ***, p < 0.001.

**Figure 4.**
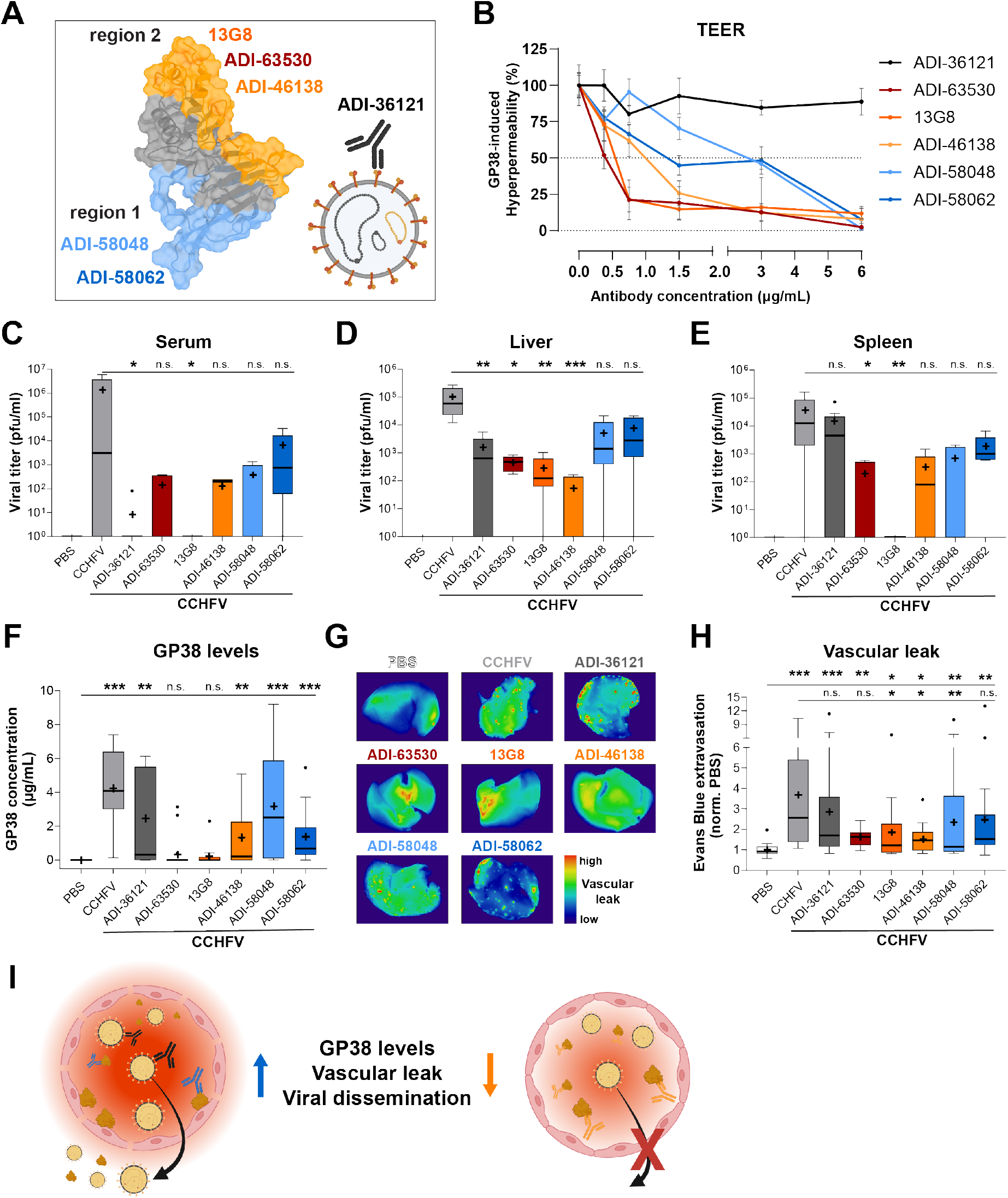
GP38-specific, non-neutralizing antibodies protect against endothelial barrier dysfunction and vascular leak. (A) Surface representation of GP38 from CCHFV strain IbAr10200 (PDB ID: 6VKF) colored by GP38-specific mAbs to antigenic region 1 (blue) and antigenic region 2 (orange). Schematic of virion with Gc-binding mAb (black). (B) HPMEC were treated with GP38 complexed with varying amounts of the indicated mAb, and TEER was measured after six hours incubation. The area under the curve of the negative peaks was calculated and normalized to 100% for GP38 in the absence of mAbs (n=4). (C-E) Viral load in the serum (C), liver (D) and spleen (E) of mice treated with an anti-IFNAR mAb MAR1-5A3 three days after PBS treatment or CCHFV infection in the presence or absence of GP38 mAb treatment (n≥5). (F) Concentration of GP38 in the serum of PBS-treated or CCHFV-infected mice determined by quantitative sandwich ELISA (n≥8). (G) Vascular leak visualized by Dextran-680 in the liver of CCHFV-infected mice treated with and GP38-specific mAbs as indicated. One representative image is shown. (H) Quantification of extravasated tracer dye Evans Blue in the liver of CCHFV-infected and GP38 mAb-treated mice (n≥13). (I) Visualization of proposed mode of action of protective, non-neutralizing GP38-specific mAbs in preventing GP38-mediated vascular leak. Data shown as a Tukey distribution with the median as a bar and mean as +. Statistical comparisons were performed by Kruskal-Wallis test with an uncorrected Dunn’s test with n.s., non-significant, p > 0.05; *, p < 0.05; **, p < 0.01; ***, p < 0.001.

The observation that recombinant GP38 alone can trigger endothelial barrier dysfunction led us to further hypothesize that GP38 might disrupt the EGL, as previously shown for flavivirus NS1 *(22)*. The EGL is a thick layer of glycosaminoglycans and proteoglycans covering the apical side of the endothelium that is thought to protect the endothelial cells from the shear stress of blood flow. Measurement of the integrity of EGL components sialic acid, heparan sulfate and chondroitin sulfate revealed significant reduction in the amount of sialic acid and chondroitin sulfate, but not heparan sulfate, on the surface of HPMEC after GP38 treatment (Fig. 3G-H). We also measured disruption of cell-surface sialic acid in skin- and liver-derived endothelial cell lines (HMEC and human hepatic synovial endothelial cells HHSEC, respectively) upon GP38 treatment (fig. S4C-F). Thus, our findings support the hypothesis that the observed endothelial dysfunction triggered by GP38 is induced through an endothelial cell-intrinsic pathway, leading to disruption of EGL components and endothelial hyperpermeability.

### GP38-specific, non-neutralizing antibodies protect against endothelial barrier dysfunction and vascular leak

After establishing the capacity of GP38 to trigger vascular leak independently of viral infection, we investigated if protective but non-neutralizing GP38-binding antibodies could act in part by blocking the interaction of GP38 with endothelial cells, thus inhibiting GP38-triggered endothelial hyperpermeability and thereby reducing vascular leak and viral dissemination. To test this hypothesis, we utilized a panel of previously characterized GP38-specific mAbs that showed differences in their protective capacity depending on their binding epitopes *(9)*. The epitopes of these mAbs were binned into two predominant antigenic regions, binding across the variable loop 2 and C-terminal β-hairpin (overlapping bins I and II in region 1, shades of blue) or binding across the 3-helix bundle and the loops connecting the β-sandwich (overlapping bins III through V in region 2, shades of orange), with mAbs binding to region 2 possessing a greater cross-clade protective efficacy against lethal challenge in two mouse models of CCHFV infection *(9)*. In addition to GP38-specific mAbs, the Gc-binding protective neutralizing antibody ADI-36121 *(7)* was included to distinguish GP38-specific observations from overall reductions in viral load (Fig. 4A).

We evaluated the capacity of this well-characterized panel of mAbs to block GP38-induced endothelial hyperpermeability by determining the IC_50_ of each mAb complexed with GP38 in the TEER assay. As expected, we found that the Gc-binding mAb did not block the induction of endothelial barrier hyperpermeability by GP38 (Fig. 4B, black). In contrast, we found that all GP38-specific mAbs could reduce endothelial hyperpermeability, with region 2 antibodies (orange) exhibiting an approximately 10-fold higher protective capacity than region 1 antibodies (blue) (Fig. 4B, fig. S5A-F), correlating with their protective profile during authentic CCHFV infection *in vivo (9)*.

To test if this possible mode of action is relevant during viral infection *in vivo*, we again used our vascular leak model. Here, mice were infected with CCHFV (IbAr10200), transiently immunosuppressed with mAb MAR1-5A3, and treated with 0.2 mg of the GP38 mAb panel as well as the Gc-binding, neutralizing antibody (ADI-36121) 30 minutes post-challenge. At the height of disease (3 days post-challenge), tracer dyes were injected intravenously, and serum and organs (liver, spleen, and kidney) were collected to assess tissue-specific viral load by plaque assay, circulating GP38 levels by ELISA, and induction of vascular leak by measuring dye extravasation. We found that the neutralizing Gc-specific mAb had the strongest effect on serum viral load, whereas region 2 GP38-specific mAbs exhibited the strongest reduction of viral load in the distal organs, including the liver, spleen and kidney – indicating that these mAbs limit viral dissemination (Fig. 4C-E, fig. S6A). While the region 1 GP38-specific mAbs also reduced overall viral load in both serum and distal tissues, they were less effective than region 2 mAbs.

Finally, we hypothesized that the observed reduction in viral load in distal tissues might be due to a reduction in circulating GP38 levels and resulting decrease in vascular leak. We determined the levels of circulating GP38 in the serum by ELISA and found that region 2 GP38-binding mAbs strongly reduced levels of GP38 in the bloodstream, while region 1 GP38 and Gc-specific mAbs only moderately decreased GP38 levels (Fig. 4F). Next, we investigated the levels of vascular leak in the liver using our systemic leak model. As expected, we observed a significant increase in vascular leak in CCHFV-infected mice compared to the PBS control group (Fig. 4G-H, fig. S6B, 7). Interestingly, while we found a reduction in viral load and GP38 levels with the Gc-specific mAb, we observed similar levels of vascular leak as CCHFV-infected animals not treated with mAbs (Fig. 4G-H, fig. S6B, dark gray bars). In contrast, the more protective region 2 GP38-specific mAbs strongly reduced the occurrence of foci of vascular leak and overall amounts of extravasated dye in the liver of CCHFV-infected animals. While region 1 GP38-specific mAbs were able to reduce vascular leak in the distal organs, this was less significant than the reduction observed for region 2 mAbs.

In summary, GP38-specific non-neutralizing mAbs were able to reduce circulating GP38 levels, vascular leak, and viral dissemination into distal organs (Fig. 4I). Importantly, the GP38-specific mAbs shown to be the most protective against lethal challenge (region 2 mAbs) *(9)* were also the most potent at reducing circulating GP38 levels, diminishing vascular leak and curtailing viral dissemination to distal organs. The moderate effect of the virus-neutralizing anti-Gc mAb indicates that the observed reduction of vascular leak upon treatment with GP38-binding mAbs is not only a result of overall reduction in viral load but rather is specific to a reduction in circulating levels of GP38 and direct blockade of the function of GP38 in inducing vascular leak. The differences between region 1 and region 2 GP38-specific mAbs suggests that epitopes within the N-terminal region comprising the 3-helix bundle and loops of the β-sandwich on GP38 are important structural targets to prevent viral dissemination and GP38-triggered endothelial dysfunction.

## Discussion

In this study, we describe the induction of endothelial barrier dysfunction and vascular leak during CCHFV infection triggered by the secreted glycoprotein GP38 through interaction with the endothelial cell barrier. We propose that the observed barrier dysfunction benefits the virus by providing access to susceptible target cells in distal tissues, including the liver and spleen, for viral dissemination and amplification. We further show that GP38-triggered vascular leak can be abrogated by antibodies targeting specific regions of GP38, identifying a novel mode of action of these protective, non-neutralizing antibodies. Fc-dependent and complement-dependent clearance of infected cells potentially contributes to the observed protection, although these mechanisms do not fully explain or correlate with the protective efficacy of GP38-specific mAbs *(13, 14)*. Furthermore, the association of reduction in GP38 levels with protection against the induction of vascular leak suggests a direct role for extracellular GP38. Therefore, we propose that antibody binding of GP38 prevents its interaction with endothelial cells resulting in reduction of vascular leak, and that this is an additional mode of action of the non-neutralizing GP38-specific mAbs that contributes to protection and aligns well with the observed protective efficacy.

The newly established animal model of vascular leak for CCHFV and testing of antibodies for their capacity to block endothelial hyperpermeability will be highly relevant for future evaluations of vaccine and therapeutic antibody candidates. The quantitative GP38 ELISA we developed has potential to measure GP38 levels as a biomarker of endothelial dysfunction during CCHFV infection as a potential method for clinicians to identify patients who are at high risk of progression to severe CCHF, as has been previously proposed with GP38 and DENV NS1 *(13, 27)*. This study was limited to the IbAr10200 strain, necessitating future studies to analyze potential differences between CCHFV isolates and their respective GP38 proteins, considering the substantial genetic diversity of CCHFV strains *(9, 28, 29)*. Understanding differences in vascular leak induction across CCHFV clades is particularly important for the development of potent, broadly effective CCHFV vaccines and therapeutics.

The relative contribution to overall CCHFV pathogenesis of the endothelial cell-intrinsic pathway triggering vascular leak that we describe remains to be determined. Other factors, including the release of pro-inflammatory cytokines, likely play a role in exacerbating endothelial barrier dysfunction, especially in later stages of infection. In future studies, identification of residues critical for GP38-mediated endothelial dysfunction and subsequent mutagenesis in a reverse genetic system is needed to test the importance of this pathway in overall viral pathogenesis. Further, our study opens a unique avenue to investigate host factors and pathways involved in GP38-mediated endothelial dysfunction that could provide new targets for therapeutic interventions. In summary, this study provides the first evidence that GP38 is a CCHFV virulence factor or viral toxin with a direct role in viral pathogenesis that is independent of its function in viral glycoprotein biogenesis. Our findings emphasize the importance of including GP38 in CCHFV vaccine design and provides a starting point for the rational design of GP38-targeting anti-CCHF therapeutics.

## Material and Methods

### Study design

We designed this study to test the capacity of CCHFV GP38 to trigger endothelial barrier dysfunction and the ability of GP38-specific non-neutralizing antibodies to inhibit this pathogenic mechanism, thereby conferring protection from lethal CCHFV infection in mouse models. We established an assay for the quantification of vascular leak in a mouse model of CCHFV disease, using PBS as a negative control. The quantitative GP38 ELISA was first tested using a dilution series of recombinant GP38 to determine the linear range, and mouse serum samples were diluted accordingly. We used ovalbumin as a negative control in comparison to GP38-injected mice in systemic vascular leak assays, and for localized dermal leak assays, PBS or DENV NS1 were used as negative or positive or controls, respectively. All mouse experiments were performed in two separate experiments with at least 4 mice per group. For the investigation of endothelial barrier function *in vitro*, we tested recombinant GP38 alongside positive controls (DENV NS1, TNF α) and negative controls (PBS/untreated cells, CCHFV Gc protein) and performed each experiment at least three times unless otherwise indicated in the figure legends. The same controls were used for measurement of the integrity of endothelial glycocalyx layer components, and two fields of view were acquired per condition, with the mean value plotted as the biological replicate. Exclusion criteria were determined before experiments were performed and included (i) failure of positive controls to induce an >1.5-fold increase in vascular leak above the negative controls in the localized dermal leak assays and (ii) differences in TEER values at baseline exceeding 20%. Experimenters were not blinded during experiments, and sample size was decided based on previous experience with the performed analyses.

### Cell lines

VeroE6, an immortalized epithelial cell line isolated from the kidney of an adult female African grivet monkey (RRID: CVCL-0574) and SW-13 cells, a cell line isolated from the adrenal gland and cortex of a 55-year-old female patient with carcinoma (RRIDD: CCL-105) were obtained from the American Type Culture Collection (ATCC). Cells were cultured in Dulbecco’s Modified Eagle Medium (DMEM; ThermoFisher Scientific) with glutamine and enriched with 10% heat-inactivated fetal bovine serum (ΔFBS; Gibco) and 1% penicillin-streptomycin (ThermoFisher Scientific). Cells were maintained in a 37°C incubator supplied with 5% CO2. Cell lines were not authenticated following purchase.

Human pulmonary microvascular endothelial cells (HPMEC) [line HpMEC-ST1.6 R] were a gift from Dr. J.C. Kirkpatrick at Johannes Gutenberg University, Germany. These cells were isolated from an adult human male donor, and an immortalized clone was selected that displayed all major phenotypic makers of pulmonary endothelial cells *(30)*. The human dermal microvascular endothelial cell line HMEC-1 (isolated from a newborn male) was kindly donated by Dr. Matthew Welch, University of California, Berkeley. Human hepatic synovial endothelial cells (HHSEC) were purchased from ScienCell Research Laboratories. All endothelial cells were cultured in endothelial cell growth basal medium 2 supplemented with an Endothelial Cell Growth Medium-2 (EGM-2) microvascular cells supplemental bullet kit (Lonza) and maintained at 37 °C with 5% CO2.

### Ethics statement

Murine challenge studies were conducted under Institutional Animal Care and Use Committee (IACUC)-approved protocols in compliance with the Animal Welfare Act, PHS Policy, and other applicable federal statutes and regulations. USAMRIID is accredited by the Association for Assessment and Accreditation of Laboratory Animal Care, International (AAALAC) and adheres to the principles stated in the Guide for the Care and Use of Laboratory Animals, National Research Council, 2013. Mice determined to be moribund, in accordance with the USAMRIID IACUC approved criteria, were promptly euthanized. All experiments and procedures conducted at UC Berkeley were pre-approved by the UC Berkeley Animal Care and Use Committee, Protocol AUP-2014-08-6638-3 and conducted in compliance with Federal and University regulations. Six to ten-week-old wild-type C57BL/6J female mice were purchased from Jackson Laboratory (strain 000664) and housed under specific pathogen-free conditions at USAMRIID or the UC Berkeley Animal Facility.

### GP38 quantitative ELISA

High-binding half-area plates (Greiner Bio-One) were coated with 25µL of ADI-58048 (Mapp Biopharmaceuticals)*(9)* at 5 µg/mL. Plates were incubated overnight at 4°C. Approximately 18 hours later, plates were blocked with 150 µL/well of blocking buffer (5% milk in 1X phosphate-buffered saline [PBS] with 0.05% Tween-20; PBST) for two hours at ambient temperature. The serum was diluted 1:10 and then serially diluted 3-fold (final dilution was 1:7290) in blocking buffer. A standard curve was generated by diluting recombinant GP38 antigen (Native Antigen Co, United Kingdom) to 1000 ng/mL for the initial dilution followed by six 2-fold dilutions (final concentration was 15.625 ng/ml). After blocking, liquid was removed by flicking, and 25 µL of each dilution (samples and standards) was added to plates in duplicate and incubated for 2 hours at ambient temperature. Plates were washed 3X with PBST and then incubated with 25 µL/well of biotinylated ADI-63530 (Mapp Biopharmaceuticals)9 diluted 1:5000 in blocking buffer for one hour at ambient temperature. Plates were washed 3X with PBST and then incubated with 25 µL/well of horseradish peroxidase (HRP)-conjugated streptavidin diluted 1:10,000 in blocking buffer for one hour at ambient temperature. Plates were washed 3X with PBST and incubated with 25 µL/well TMB substrate (ThermoFisher Scientific) for 30 minutes at ambient temperature. Plates were read at 450 nm (SpectroMax M5; Molecular Devices) after the addition of 25 µL/well stop solution (0.16M sulfuric acid).

### *In vivo* challenges

Eight-to 10-week-old male and female C57BL/6J mice (strain #000664; The Jackson Laboratory) were exposed intraperitoneally (IP) to 100 PFU of CCHFV-IbAr10200 or an equal volume (200 µL) of PBS vehicle control. Mice were treated IP with 0.2 mg of the indicated mAb or an equivalent volume (200 µL) of PBS vehicle 30 minutes post-challenge. Twenty-four hours post-challenge, mice were transiently immunosuppressed via treatment IP with 2 mg/mouse of anti-mouse IFNAR-1 antibody (clone MAR1-5A3; PLATINUM Functional grade; Leinco Technologies). Seventy-two hours post-challenge, mice were given a cocktail of a 500 g/ml stock of dextran conjugated to AlexaFluor 680 (50 µL; D34680; ThermoFisher Scientific) and 2.5% weight/volume (w/v) Evans Blue (10 µL; Sigma; cocktail delivered in 60 µL total volume) via intravenous tail vein injection. The dye was allowed to circulate for 4 hours, at which point mice were euthanized and whole blood and liver were collected. The liver was cut in half and each half was placed in a separate container in 10% neutral buffered formalin solution (Valtech Diagnostics Inc.). For the first set, liver tissue was fixed overnight and approximately 12-24 hours post-fixation, fluorescence signal accumulation was visualized using a Chemidoc imaging system (Bio-Rad) at a wavelength of 680 nm. Leakage was quantified using ImageJ software and reported relative to PBS controls. For the second set, liver tissue was moved to a fresh tube with 1 ml of 10% neutral buffered formalin solution and incubated at 60°C for 24 hours for Evans Blue dye extraction. After 24 hours, samples were centrifuged at 13,000 × g for 10 minutes, and absorbance of supernatants was read at 611 nm (SpectroMax M5; Molecular Devices). Livers were weighed and absorbance corrected based on weight and reported relative to PBS controls. For viral load assessment, at 3 days post-challenge, mice were euthanized and tissues (liver, spleen, kidney) and whole blood were harvested for viral load assessment. Whole blood was centrifuged at 12,000 × g for 10 minutes and serum was aliquoted to a fresh tube for further analysis. Tissues were homogenized in DMEM supplemented with 5% ΔFBS, 2mM L-glutamine, and 1% penicillin-streptomycin in gentleMACS™ M tubes (Miltenyi Biotec) to generate a 10% tissue homogenate.

### Plaque assay

Serial dilutions of serum and tissue homogenates were prepared in DMEM supplemented with 5% ΔFBS, 2mM L-glutamine, and 1% penicillin-streptomycin. Following dilutions, 200 µL from each dilution was inoculated onto SW-13 cell monolayers in 6-well plates. After adsorption for 1 hour at 37°C, cell monolayers were overlaid with a mixture of 1 part 1.2% agarose (Seakem ME) and 1-part 2x Eagle basal medium, 30mM HEPES buffer, and 5% ΔFBS. Plates were incubated for 4 days at 37°C, and after 4 days, a second overlay, supplemented with 5% neutral red, was added. Plaques formed by infectious CCHFV-IbAr10200 were then counted 6-12 hours later, and titers are shown as PFU/mL.

### Infection-independent vascular leak assays *in vivo*

Six-week-old C57BL/6J female mice were anesthetized and intravenously injected with 0.1 mg OVA or CCHFV GP38 (IbAr10200) via retroorbital administration. Protein was allowed to circulate for 48 hours, and mice were observed daily for behavioral changes. Evans Blue (50 µL, 0.5%) was intravenously injected via the retro-orbital route. Three to four hours post-treatment, mice were euthanized, and organs extracted, and organs were transferred into 10 mL of neutral-buffered 10% formalin (Sigma). Organs were fixed overnight before transfer to a fresh tube containing 1 mL of formamide (Sigma). Organs were incubated for 48 hours at 60°C and then spun at 16,000 x g for 10 minutes. Supernatant was transferred into a 96-well plate and absorbance measured at 620 nm using a spectrophotometer. Relative levels of extravasated dye were calculated by normalization to the OVA-treated group. The murine dermal leak model to study the effect of GP38 on a microvascular level was performed as previously described21,24. Briefly, the dermis of 6-8-week-old C57BL/6J female mice were shaved and depilated with Nair. Three days later, GP38 alongside positive and negative controls (DENV NS1 and PBS, respectively) was intradermally injected into distinct spots (50 µL per spot) followed by intravenous injection of 10-kDa dextran conjugated to Alexa Fluor 680 (Sigma; 150 µL, 167 ng/µL). After circulation for 3 hours, the dorsal dermis was removed, and extravasated tracer dye was measured using a fluorescence scanner (LI-COR Odyssey CLx Imaging system). The mean fluorescence intensity (MFI) was measured in a consistent circular area surrounding the injection spot using Image Studio software (LI-COR Biosciences). Data points were excluded when the positive control did not lead to >1.5-fold increase in MFI.

### Transendothelial electrical resistance assay

Endothelial hyperpermeability was measured as previously described22. In brief, 50,000 cells were seeded in the apical chamber of 24-well Transwell polycarbonate membrane inserts (Transwell permeable support, 0.4 µm, 6.5 mm insert, Corning), and the lower chamber was filled with 1.5 mL of EGM2 medium (Lonza). Half of the medium volume of both chambers was changed every day for 3 days until a confluent monolayer was formed. On the day of the experiment, TEER was measured in ohms (Ω) at baseline, and then the indicated treatments were added to the apical chamber. Electrical resistance was measured at the indicated time points with an Epithelial Volt Ohm Meter (EVOM) in combination with a “chopstick” STX2 electrode (World Precision Instruments). Untreated cells and a chamber without cells were used as controls to calculate relative resistance values and subtract background, respectively. Wells with a 20% difference to the untreated control well at baseline or 24 hours post-treatment were excluded. For mAb blockade experiments, GP38 was complexed with the different mAbs at the indicated concentrations for 30 minutes at 37°C before treatment of the HPMEC.

### Endothelial glycocalyx layer assay

Endothelial cells were seeded on gelatin-coated (0.2% v/v in PBS; Sigma) coverslips at a density of 50,000 cells, and medium was changed daily for three days. On the day of the experiment, the medium was changed, and treatments added. For sialic acid staining, cells were treated with wheat germ agglutinin conjugated to Alexa Flour 647 (Thermo Fisher Scientific, W32466) at 5 hours post-treatment (hpt). For all staining conditions, cells were washed twice with PBS and fixed at 6 hpt using 4% paraformaldehyde. The other EGL components were stained using primary antibodies for heparan sulfate (amsbio, clone F58-10E6) and chondroitin sulfate (Thermo Fisher Scientific, clone CS-56).

### Expression and purification of mAbs

Variable regions were ordered as gBlocks (Integrated DNA Technologies) with a 15 base pair 5’overlap to a mouse IgKVIII secretion signal and a 15 base pair 3’overlap to the appropriate constant region (human kappa, human lambda or human IgG1). The variable regions were cloned into pCDNA 3.4 (Thermo Fisher Scientific) vectors previously constructed with a mouse IgKVIII signal sequence and each constant region. In-Fusion enzyme (Takara Bio) was used to insert the gBlocks between the secretion signal and the constant region. Antibodies were transiently expressed in ExpiCHO cells (Thermo Fisher Scientific) following the high titer protocol for CHO Expifectamine (Thermo Fisher Scientific). Cultures were spun down 9-10 days after transfection. The supernatant was filtered and loaded on to a HiTrap MabSelect SuRe affinity column (Cytiva) using an AKTA pure fast protein liquid chromatography (FPLC) system. The column was washed with 10 column volumes of PBS (pH 7.2) and antibodies were eluted with Pierce IgG elution buffer (Thermo Fisher Scientific). Fractions containing antibodies were combined and neutralized to ∼pH 7 with 1M Tris pH 7.8.

### Expression and purification of CCHFV GP38

Recombinant CCHFV IbAr10200 strain GP38 was expressed as previously reported9,14. Briefly, the pαH eukaryotic expression vector encoding IbAr10200 mucin-like domain-GP38 with a human rhinovirus 3C protease cleavage site, an 8x His tag, and a Twin-Strep tag was co-transfected with a pCDNA3.1 plasmid encoding furin at a mass ratio of 1:9 furin:GP38 to ensure cleavage of MLD from GP38. Both plasmids were transiently transfected into FreeStyle 293-F cells (Invitrogen) using polyethyleneimine. Additionally, 5 μM kifunensine was added during transfection to ensure uniform high-mannose glycosylation across GP38. GP38 was secreted into the medium, harvested, and purified using Ni-NTA resin (Thermo Scientific HisPur™ Ni-NTA Resin). Lastly, the GP38 Ni-NTA eluent was further purified, and buffer was exchanged into 1x PBS using SEC with a HiLoad 16/600 Superdex 200 column (GE Healthcare Biosciences).

### Statistical and informatic analysis

Statistical analysis and graph generation was performed using the GraphPad Prism 8 software package. Experiments were repeated at least 3 times unless indicated otherwise, and negative and positive controls were included in the experimental design for inclusion and exclusion criteria. Normality of the data was tested using the Shapiro-Wilk test, and parametric (Ordinary one-way ANOVA) or non-parametric (Mann-Whitney, Kruskal-Wallis) statistical hypothesis testing was performed accordingly, as indicated in the figure legends. The resulting p-values are shown as *, p < 0.05; **, p < 0.01; ***, p < 0.001, while non-significant differences are shown as n.s. or not indicated. Parametric data is shown as mean ± SEM, while non-parametric data is displayed as a Tukey distribution with both the median (line) and mean (+) indicated. The surface representation of GP38 was performed in ChimeraX *(31)* with the CCHFV IbAr10200 GP38 structure (PDB ID: 6VKF;14), with the antibody footprints based on the previously described epitopes *(8, 9)*. Researchers were not blinded during experiments.

## List of Supplemental Materials

Figures S1-S7

## Acknowledgements

We thank Marco Antonio Chapa and Claudia Sanchez San Martin at UC Berkeley and Kandis Cogliano, Kathleen Dempsey, and Cecilia O’Brien at USAMRIID for excellent administrative support. Confocal images were acquired with a Zeiss LSM 710 at the CRL Molecular Imaging Center at UC Berkeley, which is supported by the Gordon and Betty Moore Foundation. BioRender was used for the generation of Fig. 1A, 3A and 4I. We are grateful to Gabby Scher, Marcus P. Wong, Jaime Cardona-Ospina, Vanessa Jimenez Posada, Pedro Carneiro, Sandra Bos and P. Robert Beatty for helpful discussions and advice. We thank Eduarda Ferreira Lopes for assistance with animal husbandry.

## Author contributions

Conceptualization: FP, SRM, KC, EH

Methodology: FP, SRM, JSM, SBB, ASH, EH

Investigation: FP, SRM, XF, CKH, AW, AIK, RRB, TGB, SEL, MM, LS

Visualization: FP, SRM, XF, CKH, AIK, TGB

Funding acquisition: FP, JSM, ASH, KC, EH

Project administration: SRM, ASH, KC, EH

Supervision: SRM, DMA, JSM, ASH, KC, EH

Writing – original draft: FP, EH

Writing – review & editing: all authors.

## Data and materials availability

Materials generated in this study are available upon request.

