## Supplementary Material for "Antibodies targeting Crimean-Congo hemorrhagic fever virus GP38 limit vascular leak and viral spread"

**Figs. S1-S7**

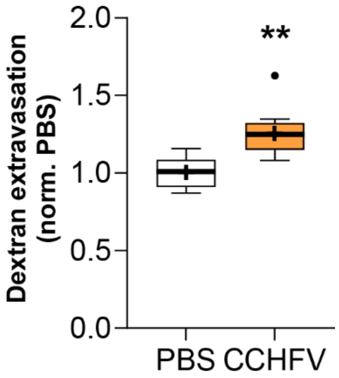


**Fig. S1. CCHFV infection leads to dextran dye extravasation.**
Extravasation of tracer dye (10-kDa dextran conjugated to Alexa Fluor 680) into the liver of PBS-treated or CCHFV-infected mice relative to PBS control animals (n=5-10). Data shown as a Tukey distribution with the median as a bar and mean as +. Statistical comparisons were performed by Mann-Whitney test with **, $p < 0.01$.

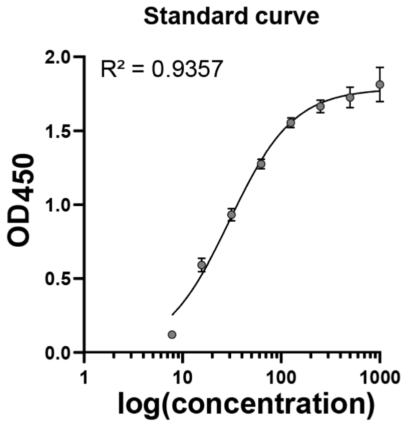


**Fig. S2. CCHFV GP38 levels can be measured by a quantitative sandwich ELISA.**
Standard curve of the quantitative CCHFV GP38 ELISA over a dilution series with recombinant GP38. Recombinant GP38 antigen (Native Antigen Co, United Kingdom) was initially diluted to 1000 ng/mL followed by six 2-fold dilutions (final concentration was 15.625 ng/ml). The captured GP38 was detected and the OD450 measured. A non-linear sigmoidal fit was used to calculate the goodness of fit, and $R^2$ is shown.

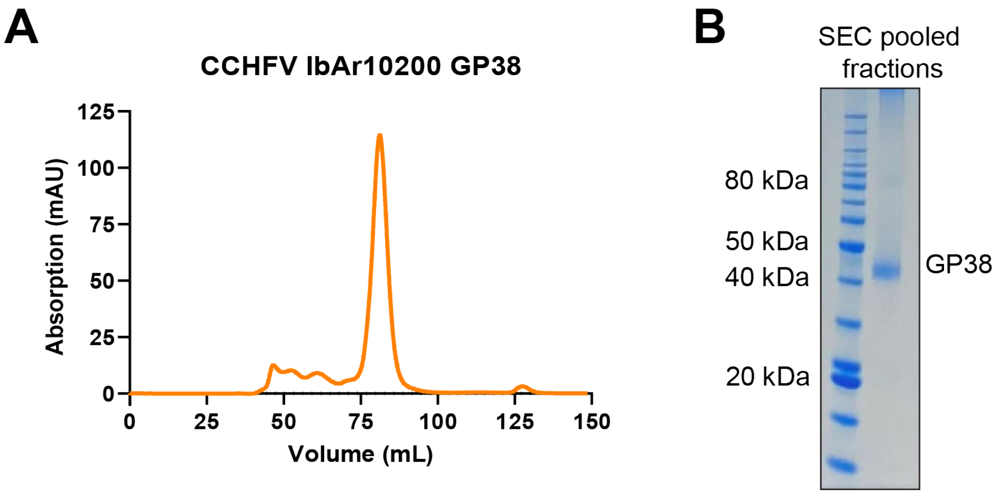


**Fig. S3. Recombinantly produced CCHFV IbAr10200 GP38 is pure and monomeric.**
(A) Size exclusion chromatography (SEC) absorption profile of recombinantly produced GP38 indicating a pure protein concentrate. (B) Coomassie staining of recombinantly produced GP38 after pooling of the GP38-containing SEC fractions. The molecular weight of the bands of the protein ladder are indicated on the left.

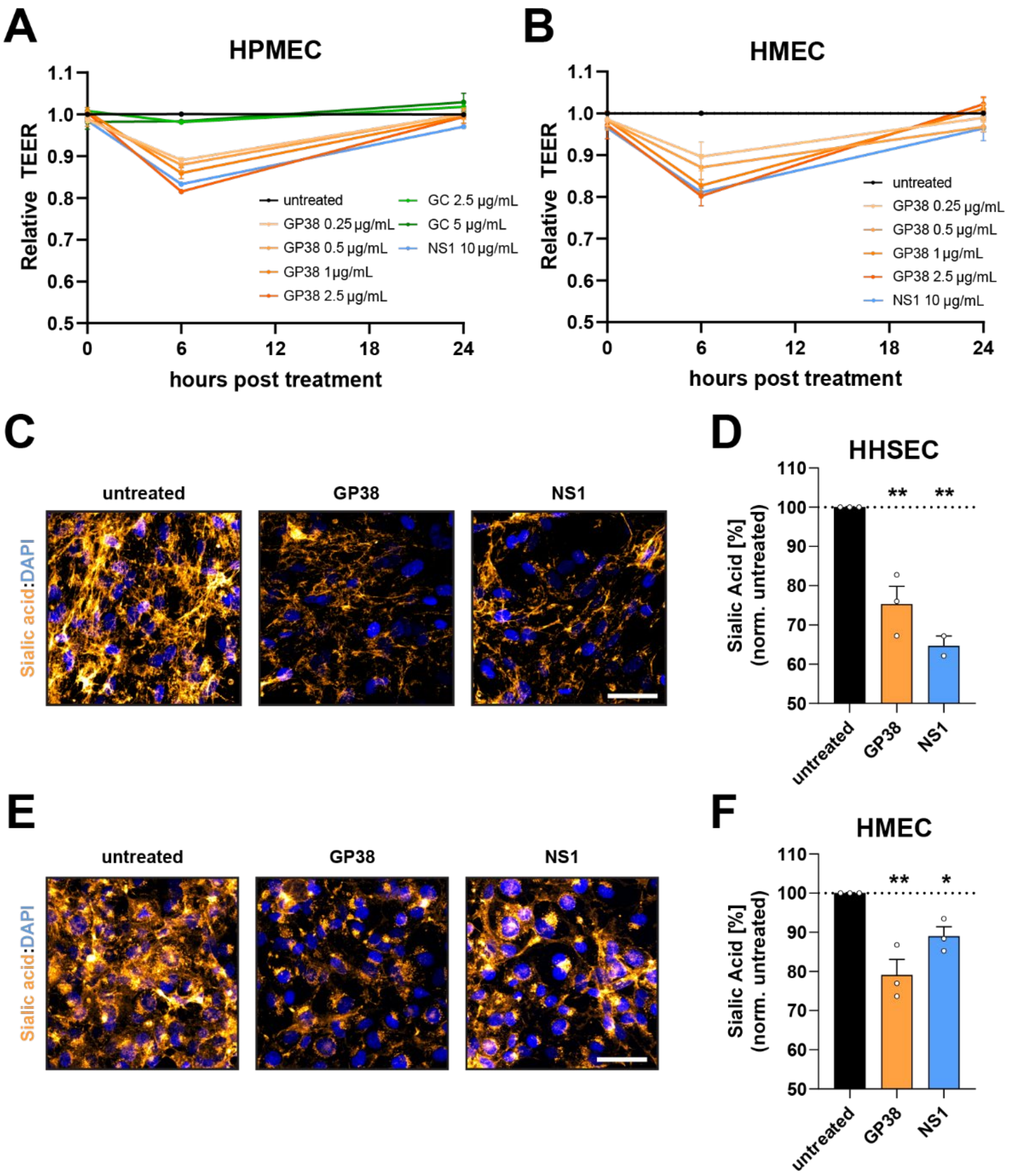


**Fig. S4. CCHFV GP38 treatment results in endothelial hyperpermeability and glycocalyx layer component disruption in lung-, skin- and liver-derived human endothelial cells.**

(A-B) Relative TEER of human pulmonary microvascular endothelial cells (HPMEC; A) or human dermal microvascular endothelial cells (HMEC; B) treated with CCHFV GP38 or left untreated and measured over a 24-hour time-course (n=3). (C-D) Human hepatic sinusoidal endothelial cells (HHSEC) were stained for EGL component sialic acid 6 hours after GP38 or dengue virus (DENV) NS1 treatment. Representative images are shown in (C) (scale bar = 50 µm), and MFI was quantified and normalized to untreated control in (D) (n≥3). (E-F) HMEC were stained for EGL component sialic acid 6 hours after GP38 or DENV NS1 treatment. Representative images are shown in (E) (scale bar = 50 µm), and MFI was quantified and normalized to untreated control in (F) (n≥3) Statistical comparisons were performed by Ordinary one-way ANOVA with Holm-Sidak test with *, $p < 0.05$; **, $p < 0.01$.

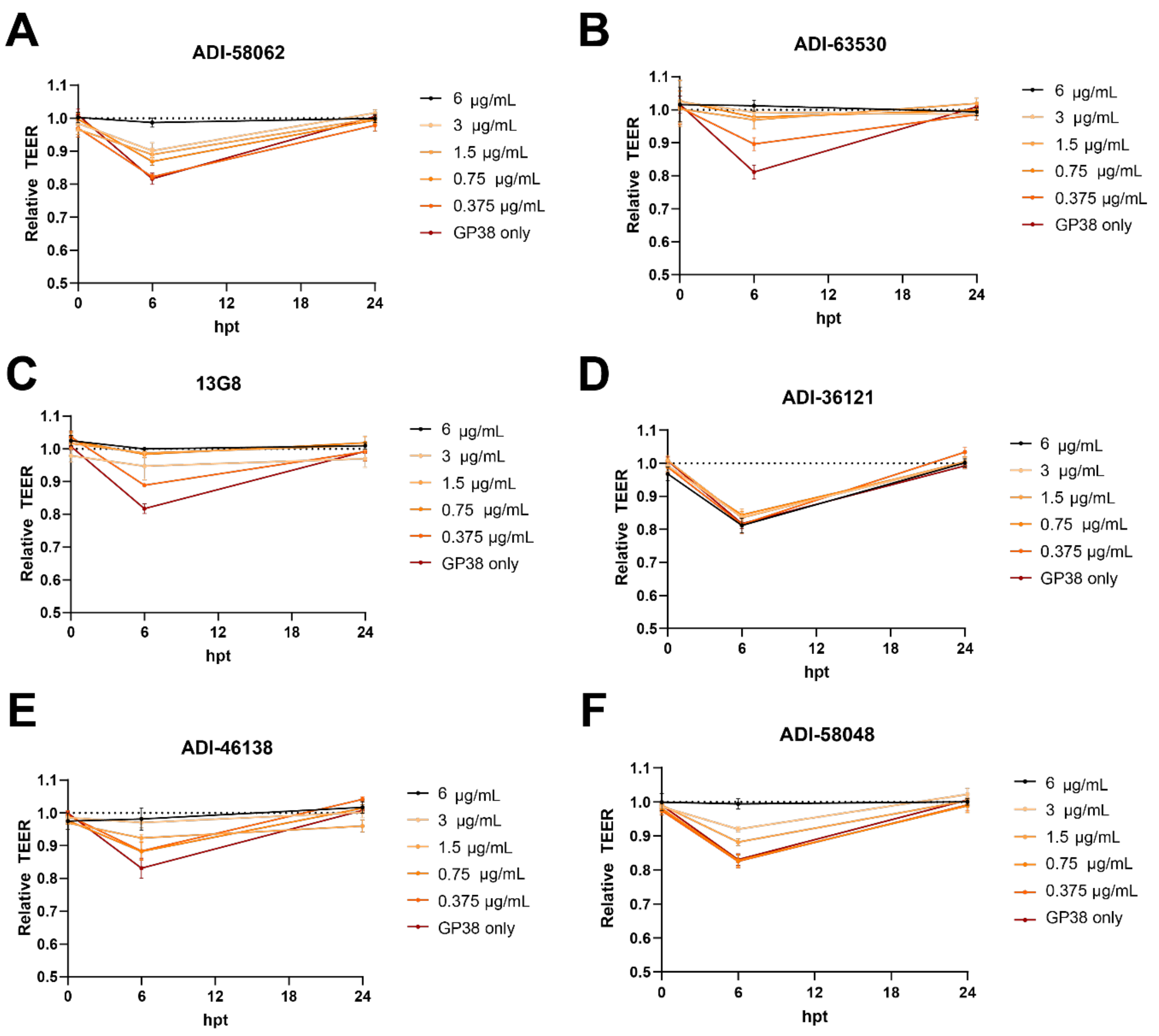


**Fig. S5. GP38-binding mAbs block endothelial hyperpermeability.**

(A-F) Relative TEER of HPMEC treated with GP38 alone or GP38 complexed with the indicated mAbs for 30 minutes at 37°C and measured over a 24-hour time-course (n=4).

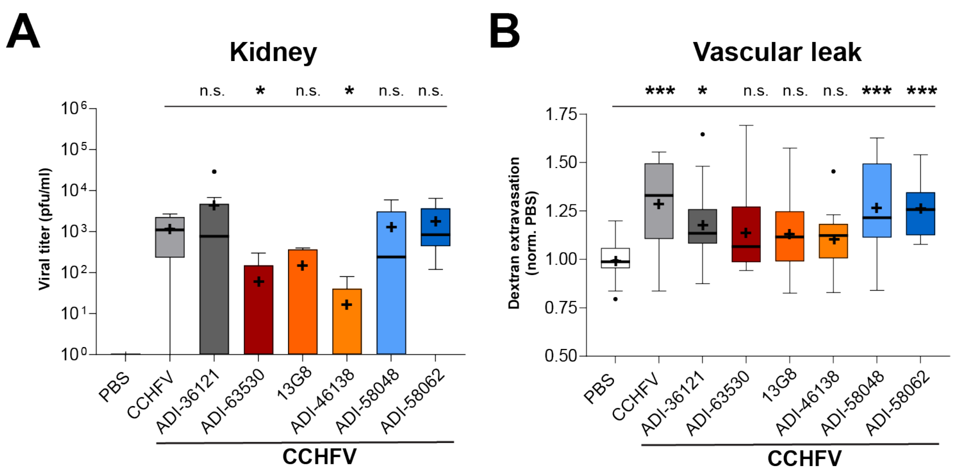


**Fig. S6. GP38-binding mAbs reduce viral load and vascular leak in distal tissues during CCHFV infection in mice.**

(A-B) Mice were infected and treated with the indicated mAbs as described in Fig. 4. Viral load in the kidney (A) of mice treated with anti-IFNAR mAb MAR1-5A3 three days after PBS-treatment or CCHFV infection in the presence or absence of GP38-specific mAb treatment ($n \geq 5$). (B) Quantification of extravasated tracer dye (10-kDa dextran conjugated to Alexa Fluor 680) in the liver of CCHFV-infected and GP38-specific mAb-treated mice ($n \geq 13$). Statistical comparisons were performed by Kruskal-Wallis test with an uncorrected Dunn's test with n.s., non-significant, $p > 0.05$; *, $p < 0.05$; **, $p < 0.01$; ***, $p < 0.001$.

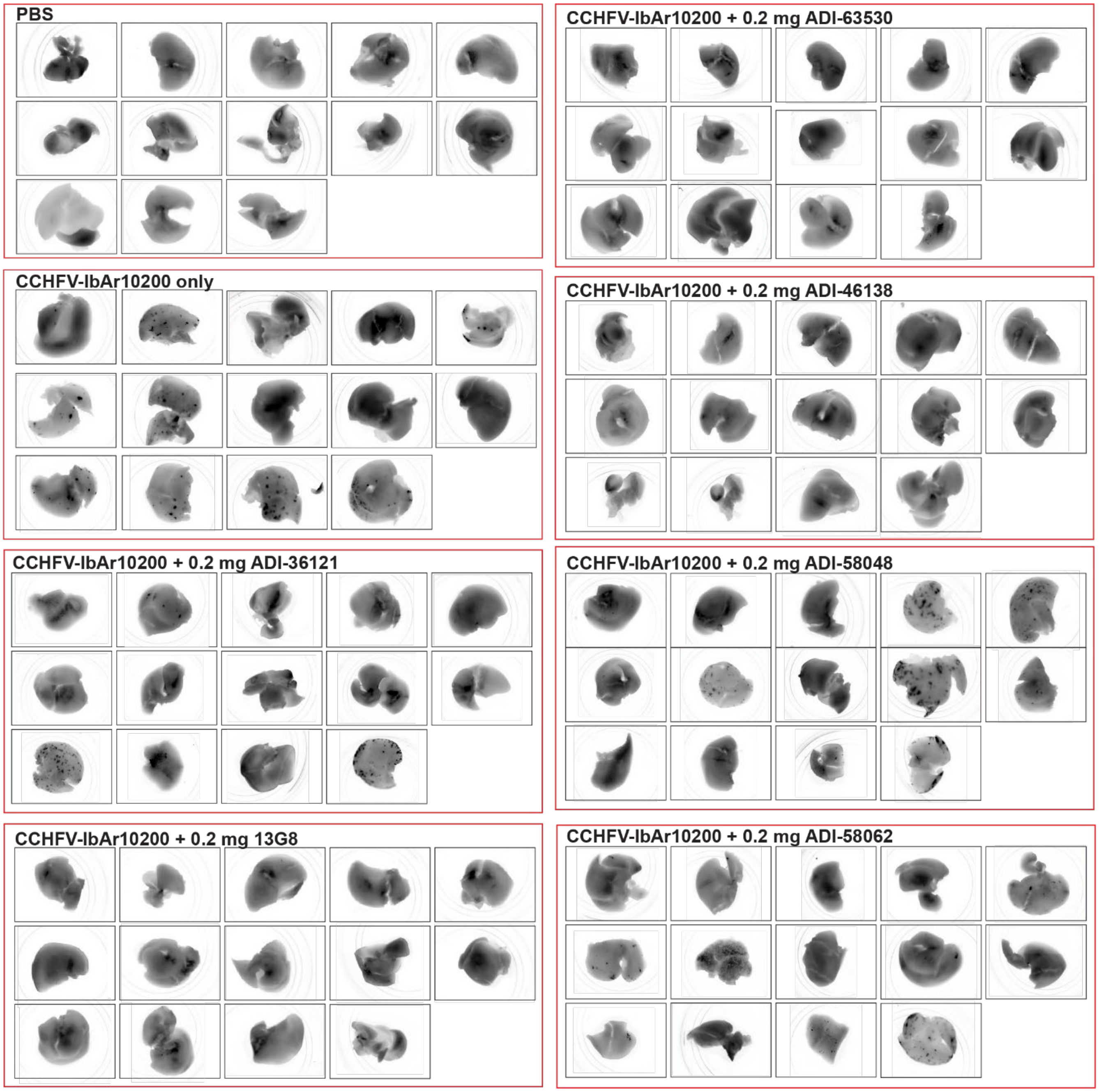


**Fig. S7. GP38-binding mAbs reduce formation of foci of vascular leak in the liver during CCHFV infection in mice.**

Mice were infected and treated with the indicated mAbs as described in Fig. 4. Livers were extracted and extravasated tracer dye (10-kDa dextran conjugated to Alexa Fluor 680) in the liver of CCHFV-infected was detected through fluorescent scanning. Foci in the liver of CCHFV-infected mice can be observed by dark spots of extravasated dye.
